# Direct evidence for pinning of single, ice-bound antifreeze proteins by subzero nanoscopy

**DOI:** 10.1101/2022.04.05.487137

**Authors:** Roderick P. Tas, Marco M. R. M. Hendrix, Ilja K. Voets

## Abstract

Ice-binding by antifreeze proteins (AFPs) reduces freezing temperatures and arrests ice-crystal ripening, making AFPs essential for survival in ice-laden environments and attractive as biocompatible antifreezes. Whilst their activity was identified over 50 years ago, the physical mechanisms are still debated because experimental insights at the molecular scale remain elusive. Here we introduce optical nanoscopy to resolve the ice/water interfacial dynamics of single AFPs. Using this method, we demonstrate pinning of individual proteins. Surprisingly, this quasi-permanent pinning is lost when freezing point depression activity is inhibited by a single mutation in the ice-binding site. These findings provide direct experimental evidence for the adsorption-inhibition paradigm, pivotal to all theoretical descriptions of activity and offer new insights in the molecular mechanisms by which these biological antifreezes function.

## Introduction

Avoiding freezing is essential for the survival of many cold-blooded species that live at subzero temperatures, such as fish, plants and bacteria. To cope, these organisms express a wide variety of structurally diverse and highly potent antifreeze proteins (AFPs) that selectively bind to ice-crystal facets to prevent rapid outgrowth of internal ice, damaging tissues and blocking circulation (*1*–*3*). AFPs function by modulating the growth of ice crystals and can lower their freezing point (thermal hysteresis, TH activity) below which outgrowth occurs explosively. Additionally, all AFPs inhibit ice recrystallization. This ice recrystallization inhibition (IRI) activity prevents many small, often harmless, crystals from ripening into large crystals that can interfere with cellular and vascular function (*2, 4*). Whilst conventional antifreezes are only effective in large quantities, AFPs function at nano- to millimolar concentrations, making them also interesting for various applications, ranging from food technology and regenerative medicine, to construction and transportation(*1, 5*–*8*).

The central framework on which all (theoretical) descriptions of AFP activity are based, is the adsorption-inhibition paradigm. Pivotal to this framework is the assumption that active AFPs lower the freezing point of ice when tightly bound; i.e., pinned, to the ice/water interface (*9*–*11*). However, the exact nature of this AFP/ice interaction and its consequences for both TH and IRI activity are still elusive. Computational studies offered a molecular perspective complementary to extensive experimental studies on the ensemble level (*12*–*17*), but direct single-molecule studies on AFPs interacting with nascent ice crystals remained hitherto out of reach. Here we present a strategy to bridge this gap and address the central paradigm for AFP activity at the single molecule level. We adopted subzero optical nanoscopy through single-molecule localization microscopy (SMLM)(*18, 19*) to directly visualize the dynamics of single AFPs at the ice/water interface. With this approach, we were able to demonstrate pinning of TH and IRI active type III antifreeze proteins from ocean pout. Surprisingly, pinning vanished when TH activity was compromised by a single point mutation in the ice-binding site of the protein, revealing that the IRI activity that remained can depend completely on transient AFP interactions with ice. These single-molecule tracking experiments demonstrate the potential of subzero nanoscopy and show how the interaction of individual antifreeze proteins with ice dictates their activity.

## Results

To directly measure the interactions of AFPs with ice, at the single molecule level, we set out to apply optical nanoscopy in temperature ranges of a few degrees Celsius below zero. An objective cooler and a dedicated sample stage were essential to achieve the precise temperature control required to generate and stabilize polycrystalline and single ice crystals with <0.1°C accuracy in the presence of fluorescently tagged AFPs **(Fig. 1A**,**B; Fig. S1)**. Our in-house built stage is inspired by a previously reported Peltier-based cooling stage(*20*) and adapted to be compatible with a Nikon dSTORM super-resolution microscope **(Fig. S1)**. Next, to achieve SMLM of single AFPs, we generated recombinant AFPs, fused to photoswitchable fluorophores (e.g. mEos3.2) for single-particle tracking photoactivated localization microscopy (sptPALM) experiments(*21*). As a result, UV induced stochastic photoconversion allows for visualization of individual AFPs at ice/water interfaces **(Fig. 1B)**. Additionally, this approach allowed visualization of the entire AFP population by (total internal reflection fluorescence) TIRF prior to sptPALM acquisitions **(Fig. 1B; Fig. S2)**. The ability to image ice crystals and visualize single, ice-bound AFPs in a high magnification optical setup enables direct measurements of the position and dynamics of AFPs with high spatiotemporal resolution for the first time **(Fig. 1C)**. The central paradigm, whether or not AFPs are pinned at the ice/water interface, underlying virtually all theoretical models of AFP activity can now be tested experimentally.

**Figure 1:**
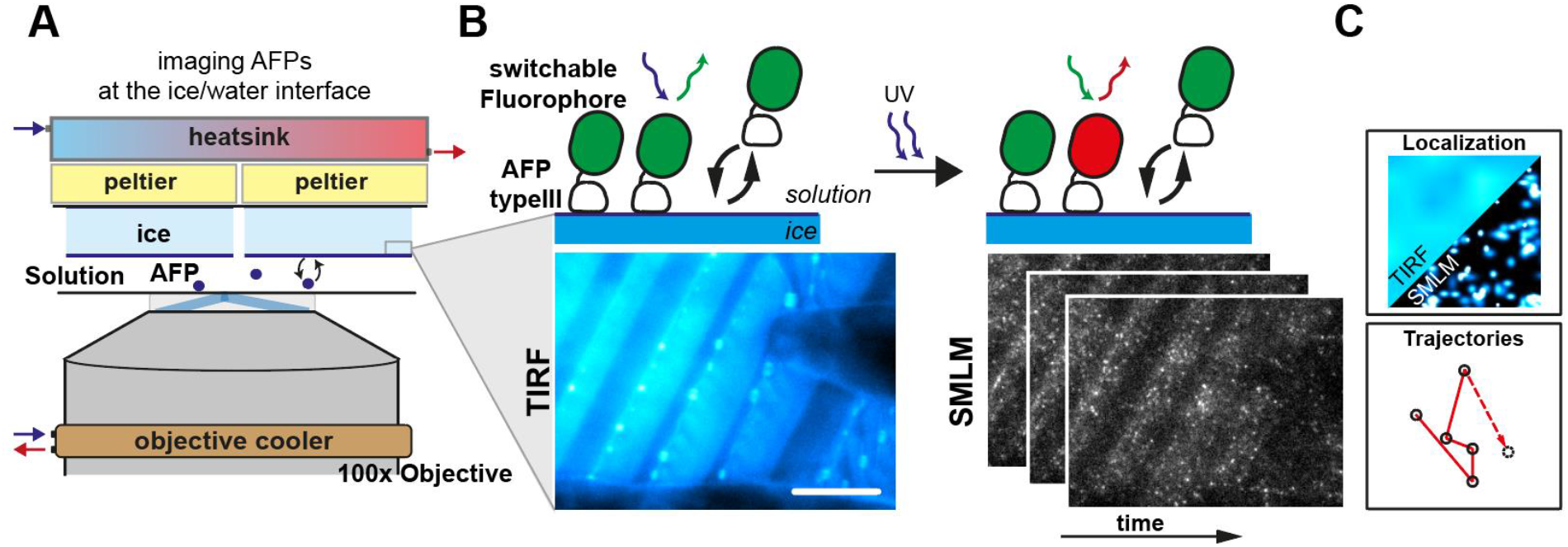
Probing single ice-binding proteins at the ice/water interface by sptPALM. A) Schematic illustration of the optical setup and generation of polycrystalline ice for sptPALM acquisitions. A stage with two Peltier elements and an objective cooler are added on an inverted TIRF microscope to control sample temperature below zero degrees Celsius. B) Schematic representation of single photoconvertible fluorophore labelled AFPs in the sample and corresponding diffraction limited TIRF image of mEos3.2-typeIII AFP on parallel crystals of polycrystalline ice. UV illumination induces stochastic photoconversion of mEos3.2 to allow single molecule detections over time. C) Single molecule localizations are used to extract the position (top) and dynamics (bottom) of AFPs at the ice/water interface with diffraction-unlimited resolution. Scale bar: 10 μm (b)

To interrogate the relation between AFP activity and dynamics at the ice/water interface, we selected the QAE isoform of the type III AFP expressed in the Ocean Pout *Z. americanus* (abbreviated as QAE). This AFP is well-characterized, easily expressed and exhibits both IRI and TH activity(*12, 22, 23*). Additionally, mutation of threonine 18 to asparagine (T18N) in its ice-binding site results in loss of TH, but not IRI(*22*–*24*), making this AFP ideally suited to study the relation between the biophysical behavior and the two modes of activity. A mNeonGreen fusion of recombinant QAE was designed for diffraction limited fluorescent measurements and a N-terminal mEos3.2 fusion for sptPALM acquisitions, both of which were purified from bacteria (**Fig. S2A)**. Activity assays showed that fusion of this small (∼7 kDa) AFP to a relatively large fluorescent protein (25 kDa) did not perturb the TH (0.6-0.8 °C at 10 mM) nor IRI (*C*_i_ ∼ 0.95 mM) activity **(Fig. S3, see also Fig. 3A)**. Fluorescently labelled mNeonGreen-QAE AFP adsorbed and accumulated selectively at primary prism planes only, shaping single ice crystals into hexagonal bipyramids consistent with its previously described binding preference(*5, 25, 26*) **(Fig. S2C-E)**.

Next, we performed sptPALM experiments on stochastically nucleated polycrystalline ice using the mEos3.2-QAE construct **(Fig. 2A-D)**. Due to its selectivity and random ice-crystal orientation in the sample, QAE did not decorate all ice/water interfaces in the field of view. Ice-planes densely covered with AFPs could be identified as the primary prism or pyramidal crystal plane **(Fig. 2B)**. Subsequent, fast sptPALM reconstructions of these planes, parallel to the imaging plane, revealed that QAE AFP localized densely and specifically at the ice/water interface **(Fig. 2C**,**D)**. Interestingly, individual AFPs in the acquisition displayed little motility and single-tracks collected for ice-bound and coverslip immobilized AFPs were essentially indistinguishable **(Fig. 2D-H; see also supplemental movie 1)**. Mean square displacement (MSD) analysis at approximately 1, 3, and 10 mM QAE showed that the distribution of the diffusion coefficients of individual tracks displayed a single peak at very low diffusion coefficients which was identical for coverslip immobilized AFPs in the control sample **(Fig. 2I, Fig. S4)**. Similarly, the average MSD curves of all conditions, regardless of concentration, are characteristic for highly confined dynamics of virtually immobile molecules **(Fig. 2J)**. These results provide the first direct evidence, at the single-molecule level, for pinning of AFPs at the ice/water interface. Additionally, these findings show that pinning is concentration-independent, suggesting that the TH and IRI active QAE binds ice in a non-cooperative manner.

**Figure 2:**
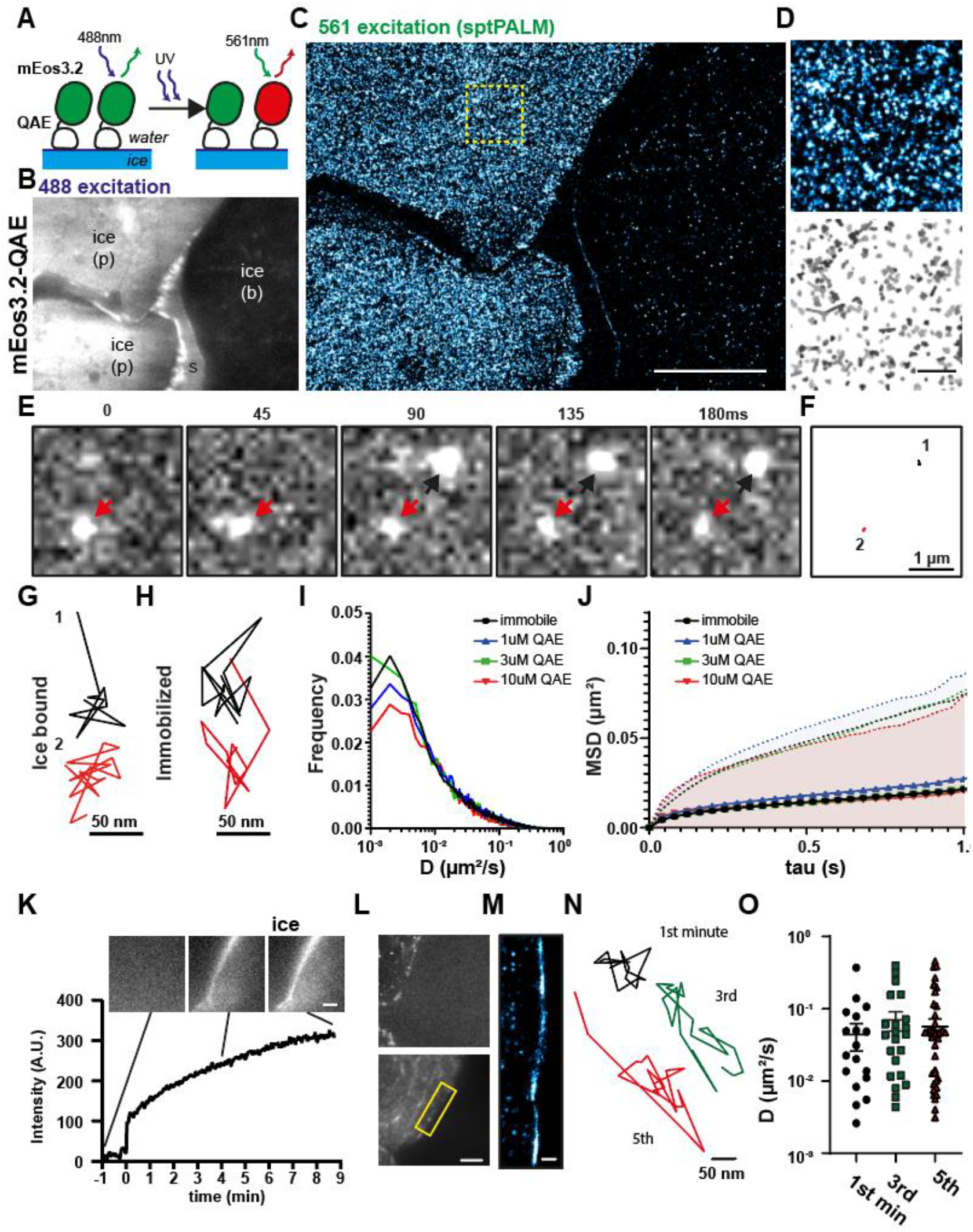
QAE AFP pins non-cooperatively at the ice/water interface. A) Schematic illustration of the sample for sptPALM acquisitions via mEos3.2-QAE on polycrystalline ice. B) Excitation of the green fluorescent channel to image selective accumulation of mEos3.2-QAE (1µM) on polycrystalline ice. Plane selective accumulation at the prism plane (p), soluble fraction (s) and non-absorbing ice planes (b) are depicted in this image. C) sptPALM reconstructions of mEos3.2-QAE on the polycrystalline ice in (b). D) Diffraction limited localizations (top) and corresponding filtered trajectories (bottom) of individual QAE molecules for the zoom indicated in (c) E) Stills from two neighboring QAE trajectories that are ice-bound. The interval between depicted frames is 45 milliseconds. The black and red arrow indicate the separate tracks. F) Diffraction unlimited reconstruction of the tracks in (e). (G,H) Highly zoomed in view of the isolated ice-bound tracks in (f) and comparable trajectories for coverslip-immobilized mEos3.2-QAE with 40ms exposure time. I) Distribution of the diffusion coefficient of coverslip immobilized QAE (Mean±S.E.M. = 0.066±0.0008 black) compared to ice-adsorbed QAE at 1 µM (0.063±0.0010, blue), 3 µM (0.069±0.0010, green) and 10 µM (0.53±0.0010, red). (N= 13851, 11910, 9122, 13393 trajectories respectively from 3 acquisition of separate polycrystalline ice assemblies) J) Average MSD curve of trajectories in (i). Error bars (dotted filled lines) represent S.D. K) TIRF acquisition to monitor accumulation of mEos3.2-AFP over time after secondary nucleation event induced by lowered temperature. Timepoint zero indicates the moment of nucleation and intensity is background corrected. Stills of the newly formed ice plane position before and after are included. L) Stills of an sptPALM acquisition before and after secondary nucleation of a new ice/water interface M) Reconstruction of the QAE afp localizations at the newly formed interface in (l) N) Example trajectories of tracks localized within the first minute after interface formation (black), the 3^rd^ (green) and after 5 minutes (red). O) Scatter plot of the diffusion coefficients (Mean±S.E.M) of trajectories observed within the first, third or fifth minute after new interface formation. Scale bar: 10 μm (b), 1 μm (d), 2 μm (k), 5 μm (l), 500 nm (m)

**Figure 3:**
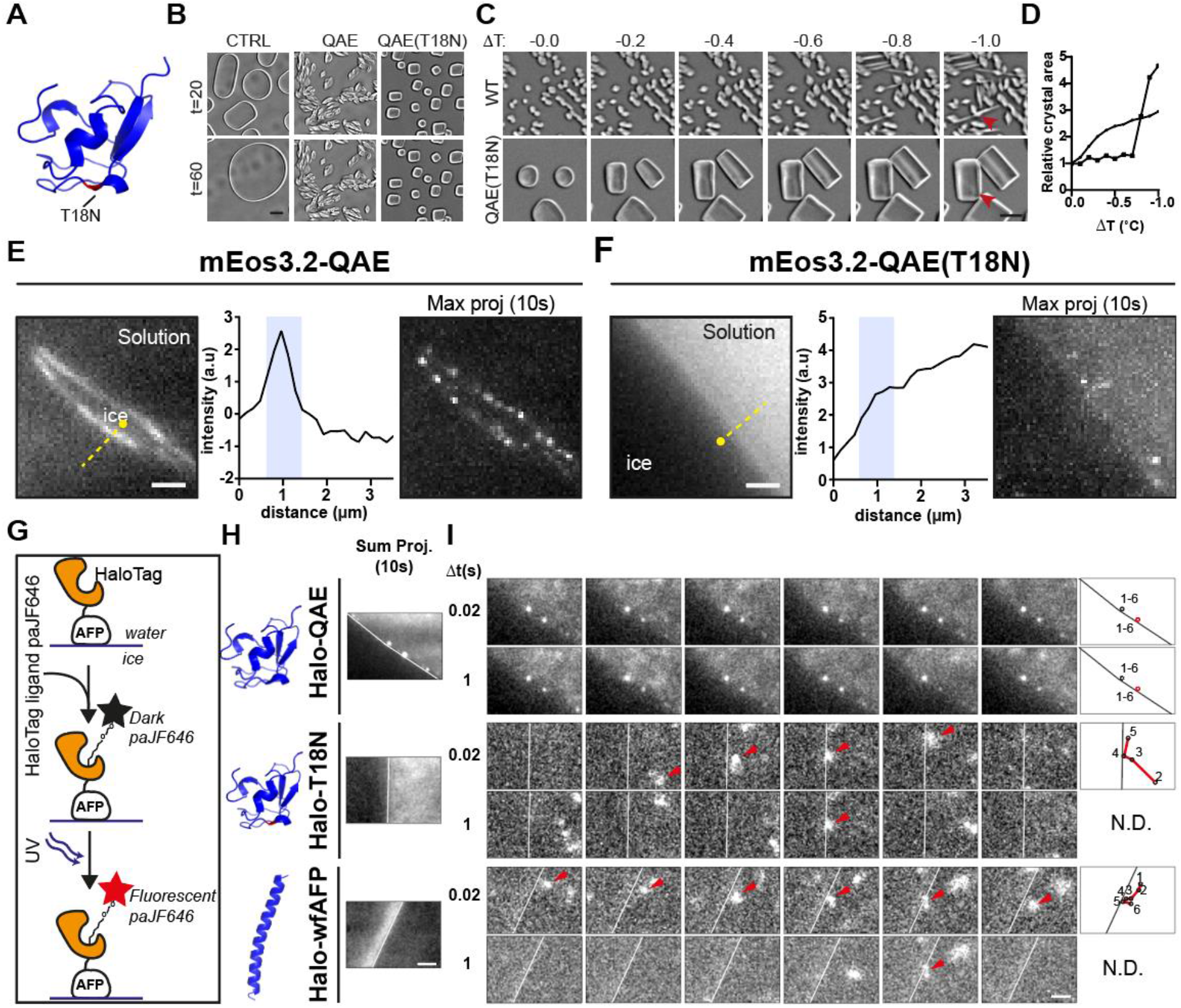
Pinning correlates with TH, but not IRI activity. A) Crystal structure of QAE (pdb: 1HG7) displaying the threonine to asparagine mutation at position 18 in red. This point mutation in the ice-binding site transforms the TH active QAE into the TH inactive T18N. B) Still images of ice crystal recrystallization assays at -7°C in 20 wt% sucrose taken after 20 and 60 minutes incubation with 10 µM QAE, 10 µM QAE(T18N) or without (CTRL). C) Thermal hysteresis activity assays of 10 µM QAE and QAE(T18N). ΔT indicates the temperature reduction from the -4°C starting point. Explosive growth of crystals in the presence of QAE occurs between 0.6-0.8°C below the melting temperature (i.e., TH = 0.6-0.8 °C), whilst crystals in the presence of 10 µM QAE(T18N) continuously grow (i.e., TH = 0 °C)). D) Relative crystal area as a function of temperature reduction for the crystals indicated in c. E,F) Diffraction limited image (left panel), intensity linescan along the yellow line (middle) and maximum intensity projection of the corresponding sptPALM acquisition (right) of ice/water interfaces in the presence of mEos3.2-QAE (E) or mEos3.2-T18N (F). The blue box in the middle panel indicates the position of the interface. G) Schematic illustration describing labelling of QAE with photo-activatable Janelia fluor646 through a HaloTag fusion. H) Sum projection for the sptPALM acquisition of Halo-tagged QAE, ; QAE(T18N and *wf*AFP labelled with paJF646. Long-term pinning events are detectable as distinct bright spots (top panel), whereas a strong diffuse signal indicates transient binding (bottom panel). I) Still images taken at 0.02 or 1 second intervals of sptPALM acquisitions of paJF646 tagged QAE, QAE(T18N) and *wf*AFP. The arrows indicate individual proteins that move along the interface. The superimposed single-molecule trajectories relative to the ice/water interface are depicted in thelast column. Tracks of T18N and *wf*AFP could be identified for short but not for long time intervals (N.D: Non-detected) Scale bar: 10 μm (b,c), 2 μm (e,f), 2 μm (h,i)

To further establish whether or not QAE-AFPs bind ice crystals in a cooperative fashion (i.e., in a density-dependent manner), QAE trajectories were acquired at variable surface coverages. To achieve this, fresh ice/water interfaces were generated through slow sample cooling, resulting in secondary nucleation events. Now, interfaces perpendicular to the imaging plane were imaged so that newly formed planes would appear in focus directly after nucleation. Monitoring AFP accumulation showed that AFP saturation did not occur until approximately 7 minutes after nucleation (**Fig. 2K**). Consequently, right after nucleation and before saturation, the interface will be decorated with increasing AFP densities. Subsequent sptPALM acquisitions showed that there were no significant differences in the dynamic behavior of single AFPs within the first minute, right after nucleation, or after accumulation for three minutes and five minutes before saturation **(Fig. 2L-O)**. In summary, these results are consistent with non-cooperative pinning of QAE AFP to ice in a quasi-permanent manner.

To unravel if both TH and IRI activity correlate with pinning, we introduced the T18N mutation in QAE. Consistent with previous work(*22*–*24*), this mutant did not exhibit TH activity whereas IRI activity was only slightly reduced compared to the wildtype construct **(Fig. 3A-D, Fig. S3, S5)**. Surprisingly, no distinct accumulation at the ice/water interface was visible for mEos3.2-QAE(T18N) by diffraction-limited fluorescence microscopy. Instead, a much more diffuse signal was observed in the soluble fraction, whereas again wild-type QAE adsorbed strongly to the ice **(Fig. 3E**,**F)**. Moreover, hardly any single molecules could be localized at the interface upon sptPALM acquisition of mEos3.2-QAE(T18N) with 40 millisecond exposure times **(Fig. 3E**,**F)**. These results suggest that pinning is required for TH activity but not essential for IRI activity. The marked difference in adsorption behavior and TH activity yet comparable IRI activity supports the premise that a weak interaction with ice is sufficient to inhibit ice recrystallization but inadequate to depress the freezing point significantly in a non-colligative manner.

To examine this interaction between AFPs and ice and its consequences for activity more closely, three AFPs were fused to a HaloTag(*27*). This tag allows coupling to the bright, stable organic photoactivatable JF646 fluorophore(*28*), to acquire single-molecule tracks with sufficient photons for single molecule localization at short exposure times (20 milliseconds) aiming to capture both pinned and more mobile proteins **(Fig. 3G)**. In addition to wildtype QAE and the T18N mutant, we included winter flounder type I AFP (pdb:1wfa) with a modest IRI activity (*C*_i_ ∼ 6 µM) and very low TH activity (TH<0.1°C at 10 mM)(*24, 29, 30*), similar to QAE(T18N). Hence, if TH does require pinning, while IRI does not, the single-molecule trajectories of *wf*AFP should resemble those of T18N and deviate significantly from the pinned QAE molecules. Indeed, as expected, pinning without any detectable mobility for as long as several seconds was visible for individual QAE’s **(Fig. 3**,**H; see also supplemental movies 2 and 3)**. In contrast, both T18N and *wf*AFP displayed highly diffusive dynamics with residence times along the interface of only one or two 20 millisecond frames. Consequentially, subsequent localizations spaced apart by short (20 millisecond) and long (1 second) time intervals could be associated with one-and-the-same AFP for the immobile QAE, but not for the mobile T18N and *wf*AFP (**Fig. 3H**,**I; see also supplemental movies 2 and 3)**. Their observed single-molecule trajectories comprised no more than a few frames separated by 20 milliseconds; i.e., at >50-fold shorter than the longest QAE tracks that could be observed using paJF646 **(Fig. 3H**,**I)**. Our results show that, transient AFP/ice interactions with residence times less than tens of milliseconds are sufficiently strong to elicit IRI activity in a non-colligative manner but not TH activity, for which pinning appears essential.

## Discussion

Internal ice formation is lethal for most living species and often detrimental to the structural and functional integrity of materials that are water-based or water-infused. The ability of AFPs to mitigate this damage by freezing point depression (TH activity) and ice recrystallization inhibition (IRI activity) is therefore highly interesting and researched, aiming to better understand the molecular mechanisms for activity on the one hand and to exploit their translational potential on the other hand(*1, 5*–*8, 31*). Here we provide the first direct experimental evidence, at the single molecule level, for quasi-permanent AFP pinning at the ice/water interface. We show that pinning is non-cooperative and correlated with TH activity. Conversely, we demonstrate that IRI active AFPs, devoid of pronounced TH activity, do not accumulate but instead bind the same interface in a highly transient fashion. The adsorption-inhibition model, which revolves around AFP pinning on ice/water interfaces, may thus rationalize TH activity, but it does not offer a satisfactory explanation for IRI activity in its current form. Therefore, future refinement aimed to establish a unifying model for the molecular mechanisms of AFP functioning should also encompass binding modes that are not quasi-permanent. Based on this work, we postulate that several binding modes may evoke IRI activity. When IRI is accompanied by TH, AFPs pin the ice/water interface and block further outgrowth and recrystallization of ice crystals in accordance with the current model. Conversely, when AFPs are exclusively IRI active, transient interaction of AFPs with ice is still sufficient to inhibit recrystallization. In conclusion, our work establishes subzero nanoscopy as a powerful tool to study the mechanisms of AFP activity at the single-molecule level. This approach has the unique potential to bridge the gap between the macroscopic observation of (arrested) ice growth and the nanoscale interactions of AFPs at the interface described in molecular dynamic simulations. This allowed us to visualize and revisit the mechanisms underlying the biological antifreeze toolbox by studying the relation between interfacial dynamics and the differential modes of AFP activity. Based on these results, existing models can be refined and new models developed to better understand how AFPs fulfill their biological roles and create bio-inspired counterparts with activities optimized and customized to function in other complex environments.

## Acknowledgements

We thank A. Aloi for initiating the construction of the cooling stage and objective collar; J. Grimm and L. Lavis at Janelia Research Campus for kindly gifting HaloLigand-paJF646 and E. Katrukha for the script to import DoM data into the MSDanalyzer Matlab class. This work is supported by the Dutch Research Council to R.P.T (NWO-VENI: 202.220, Understanding ice growth inhibition by antifreeze proteins through super-resolution microscopy), and the European Research Council to I.K.V (ERC-2014-StG: 635928).

## Supplementary Materials

Materials and Methods

Fig S1 – S5

Table S1 – S2

References (32 – 35)

Movie S1-S3

