## Supplemental information for "Direct evidence for pinning of single, ice-bound antifreeze proteins by subzero nanoscopy"

### MATERIALS AND METHODS

#### DNA constructs

First, to generate mNeonGreen-QAE (quaternary aminoethyl typeIII-AFP isoform): mNeonGreen with N-terminal NcoI; C-terminal EcoRI sites and QAE AFP with a N-terminal EcoRI; a small GAG linker; a C-terminal XhoI sites were PCR amplified with ~20 aa overhangs. Subsequently, the PCR fragments were inserted via Gibson assembly into a NcoI/XhoI restricted pET28a vector. mEos3.2-QAE, HaloTag-QAE, mEos3.2-QAE(T18N) and HaloTag-QAE(T18N) were generated by PCR amplification of the mEos, HaloTag and AFP fragments and inserted accordingly into NcoI/XhoI restricted pET28a vector. To generate Halo-wfAFP (pdb:1wfa) spaced by a small GGGGS linker and flanked by NcoI/XhoI, a gblock (IDT) was ordered, restricted and ligated in the linearized pET28a vector. All sequences were verified by sequencing. Protein sequences are depicted in table S1.

#### Protein expression and purification

The pET28a expression vectors containing the constructs for the recombinant AFP's were transformed into BL21DE3 bacteria and grown in LB containing 50 µg/mL kanamycin. At OD0.6, expression was induced by adding 1mM final concentration and cells were grown for 0.5 hours at 37 ºC, followed by overnight incubation and expression at 20 ºC. The cultures were then pelleted by centrifugation, stored on ice and the supernatant was discarded. Cells were resuspended in the appropriate buffer (QAE constructs 20mM Tris pH7.5; wfAFP construct PBS) supplemented with lysozyme and EDTA-free protease inhibitor cocktail (Sigma) and lysed by sonication on ice. The soluble fraction was separated from the insoluble fraction by centrifugation (10.000x g, 45 minutes) and incubated with washed Ni-NTA Agarose pre-charged resin (Qiagen) for 1.5 hours at 4ºC while rotating. Subsequently, the protein-loaded beads were collected in disposable columns (Bio-rad) and washed two times with 15 mL buffer, followed by two washes with buffer + 40mM Imidazole. The protein was then eluted by incubation with buffer + 400mM Imidazole. Finally, buffer exchange to the appropriate buffer for QAE or wfAFP was performed (PD-10 desalting clumns, Cytiva) and the protein was supplemented with 10% (w/v) glycerol and flash frozen in liquid nitrogen. Protein concentrations were determined by SDS-page of the purified proteins next to a BSA standard. Final concentrations after purification were 63, 99, 118, 95, 113, 111 µM for mNeonGreen-QAE, mEos3.2-QAE, mEos3.2-QAE(T18N), HaloTag-QAE, HaloTag-QAE(T18N), HaloTag-wfAFP, respectively.

#### AFP activity assays

Microscope setup: Activity assays were performed on a Nikon ECLIPSE Ci-Pol Optical Microscope under widefield illumination. Both IRI and TH activity were imaged using a Nikon L Plan 20x (NA 0.45) objective. The Linkam LTS420 stage, controlled by Linksys32 software, was used to control and stabilize the sample temperature.

Sample preparation: QAE or wfAFP were diluted in 10mM TRIS pH 7.5 or PBS respectively and supplemented with 20 w% sucrose final concentration. Prior to sample preparation coverslips (Thermo Scientific Menzel x1000, #1.5, 22x22mm) were cleaned by 5 minutes sonication in MQ followed by 5 minutes sonication in Acetone and drying under a Nitrogen gas flow. Then, a 2 µL sandwich sample was prepared between two coverslips by depositing a 2 µL sample on one coverslip and lowering the second coverslip onto it from a parallel position. The latter ensures homogeneous sample spreading.

Ice-recrystallization inhibition (IRI) assay: Samples were diluted to the indicated concentrations and frozen to -40ºC with 20 ºC/min. Samples were then heated to -10 ºC with 20 ºC/min followed by 1 ºC/min heating to -7 ºC. Ice recrystallization was then monitored by imaging at 1 minute intervals using the 20x objective.

Thermal Hysteresis (TH) measurement: TH experiments of QAE- and QAE(T18N) performed at a final concentration of 10 µM. Samples were frozen to -40ºC with 20 ºC/min. After stabilization the samples were heated to -10 ºC with the same rate and then to -4 ºC with 1 ºC/min. After stabilization of the temperature, the sample was cooled with 0.2 ºC/min and crystal growth and secondary nucleation events were monitored by imaging at 5 second intervals using the 50x Long working distance objective. A sudden increase in crystal area marks secondary nucleation at the limit of the hysteresis gap whereas continuous crystal growth in 10µM T18N indicates a lack of TH activity.

**Single molecule localization microscopy at the ice/water interface**

Optical setup: Single particle tracking PALM (sptPALM) was performed on a Nikon Eclipse Ti-E N-STORM system equipped with a Nikon 100x Apo TIRF oil immersion objective (NA 1.49) and perfect focus system. Photoactivation and excitation were performed with the 405nm, 488 nm, 561nm and 647nm excitation lasers within the MLC400B laser box (Agilent technologies) under TIRF or HiLo illumination through a quad-band polychroic mirror (Nikon 97335). An Ixon3 EMCCD (Andor) was used for detection, resulting in an effective pixel size of 160 nanometer. The microscope was fitted with a temperature controlled stage, that was adapted to fit the Nikon microscope. A pump was used to continuously flow cold water through the heat sink with 120 mL/min, to buffer the residual heat from the Peltiers. Peltiers were controlled via Meerstetter Engineering TEC controllers and software. Additionally, the objective was cooled using a fitted copper collar with a fluid channel in the center that allowed a continuous flow of ~100 mL/min of cooled water through a peristaltic pump from a cold water bath of approximately 2.5 ˚C. **(see also Figure 1a and Figure S1 for the full experimental setup).** This allowed control of the sample temperature with <0.1 degrees centigrade precision.

Sample preparation: To perform sptPALM acquisitions, 22 x 50 millimeter #1.5 coverslips (Thermo Scientific Menzel) were cleaned by five minute sonication in ultrapure water, followed by sonication in acetone and drying under nitrogen flow. Subsequently, 30-40 uL recombinant QAE or typeI AFP, diluted to the desired concentrations in 20mM TRIS pH 7.5 or PBS respectively **(see also Table S2)**, was added to the center of one coverslip and the second coverslip was applied on top to generate a thin film of sample between the coverglasses. For HaloTagged proteins 1, 0.5 or 1 µM of HaloLigand-paJF646 was added in the sample containing Halo-tagged QAE, QAE(T18N) and 1WFA respectively to achieve sparse labelling of the proteins. Samples were then mounted in the cooling stage which was placed on the Nikon dSTORM stage holder prior to imaging.

Freezing protocol: As the lower limit of the peltiers was -10 ˚C, the samples were cooled to -9.5 ˚C at a speed of ~0.5 ˚C/s upon which single or polycrystalline ice nucleated randomly throughout the sample. Samples were then slowly heated (0.05 ˚C/s) a few degrees below zero until clear ice/water interfaces could be detected and the ice-volume fraction did not exceed roughly 40 percent. Samples then stabilized and the temperature control speed was set to a maximum of 0.02 ˚C/s to make sure that the sample remained stable. Upon stabilization clear ice/water interfaces parallel or perpendicular to the imaging plane could be identified and imaged a few hundred nanometers above the coverslip. The latter ensured that aspecifically adsorbed tagged-AFPs at the coverslip did not interfere with the localizations of the specifically bound AFPs at the ice-water interfaces. Conversely, dynamics of immobilized AFPs in figure 2 were obtained by imaging of aspecifically adsorbed mEos3.2-QAE at the coverslip.

Single molecule localization microscopy (SMLM): After the temperature and sample were stabilized and interfaces were identified, SMLM was performed. First in case of mEos3.2-Tagged constructs, a diffraction limited image was acquired using the 488 lases. Then, rapid stream acquisitions were performed using the 561 or 647 excitation lasers to perform single-particle tracking photoactivated localization microscopy (sptPALM) of mEos3.2- or HaloLigand-paJF646 respectively using the imaging parameters indicated in Table S2 under HiLo illumination. During the acquisition, excitation with the 405 laser was activated at low intensity levels to stimulate photoconversion of the fluorescent molecules and maintain enough single photo-activation events to detect the antifreeze proteins throughout the acquisition. To image AFP behavior at new ice/water interfaces (Figure 2K-O). Imaging was performed at identified interfaces perpendicular the imaging plane and the sample was slowly cooled until a secondary nucleation event occurred upon which the temperature was set to stabilize.

**Analysis**

Detection and localization of sptPALM acquisitions: sptPALM acquisitions were imported in ImageJ/Fiji(*32-33*) and analyzed using the Detection of Molecules (DoM version 1.2.1, https://github.com/ekatrukha/DoM_Utrecht) plugin. To detect single molecules a signal-to-noise ratio between 3 and 4 was chosen. For single particle tracking the 'link particles to tracks' function was used with a maximum permitted distance of 2 pixels from next the detected position and a maximum linking gap of 1 frame. These settings were kept constant for all conditions so that occasionally mis-linked tracks would occur consistently and not induce differences in the mean squared displacement results. SMLM reconstructions of the AFP localizations were performed in DoM.

Diffusion analysis: Mean Squared Displacement (MSD) analysis was performed by importing the DoM linked tracks, highly similar to the approach used by Katrukha and colleages(*34*), into the 'msdanalyzer' class(*35*) for Matlab (R2021a). Tracks consisting of a minimum of 5 localization and a maximum of 30 localization were included to filter for individual mEos3.2-AFP trajectories.

Additional software: Image processing was performed using functions in ImageJ/Fiji. Additionally open source Pymol by Schrödinger, Graphpad Prism were used to visualize structures and graphs respectively. Figures were made in Adobe Illustrator.

### SUPPLEMENTAL FIGURES

**Figure S1: Temperature control stage for SMLM at the ice/water interface**

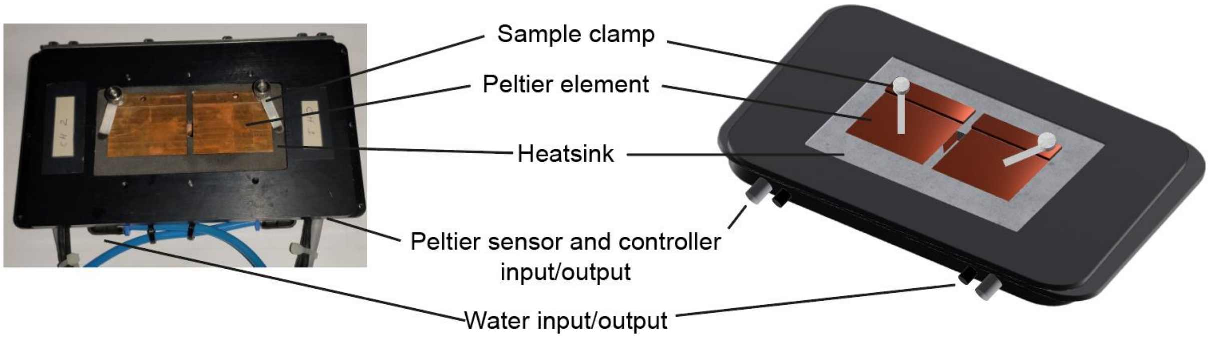


Bottom-up picture and schematic drawing of the temperature controlled stage that is used to generate ice-water interfaces on a Nikon dSTORM setup. Important elements are indicated.

**Figure S2: Selective binding of QAE AFP at ice-water interfaces on single crystals.**

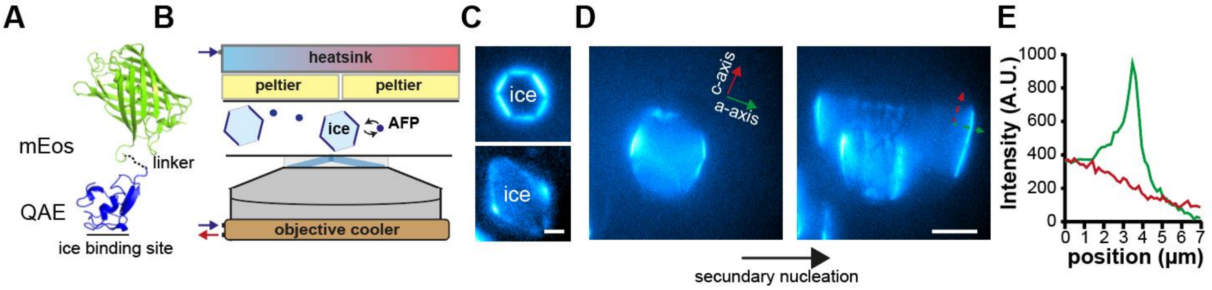


A) Structure of the mEos3.2-QAE fusion. Wild type QAE AFP is depicted in blue (pdb:1HG7), mEos (pdb:3S05) in green and the linker in black dotted line. For diffraction unlimited measurements in C-E, mEos3.2 was replaced by mNeonGreen (mNG).

B) Schematic illustration of the optical setup and sample during the generation of single ice crystals.

C) single ice crystals formed in the presence of low 5 uM and high 10 uM concentrations of mNeonGreen-QAE. mNG-QAE signal is false colored in cyan.

D) Single ice crystal decorated with QAE at the a-axis before and after secondary nucleation induced by gradual temperature reduction.

E) Background corrected intensity profile of mNeonGreen-QAE along the a-axis (green) and c-axis (red) for the lines depicted in D.

Scale bar: 2 µm (c), 10 µm (d)

**Figure S3: IRI activity of QAE AFP is not perturbed upon fusion to a fluorescent protein**

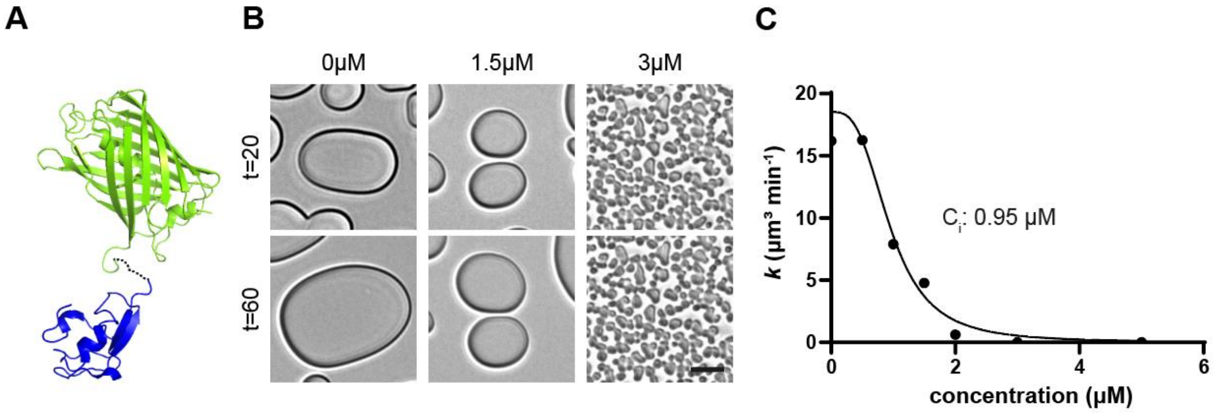


A) Structure of the mEos3.2-QAE fusion. Wild type QAE AFP is depicted in blue, mEos in green and the linker in black dotted line.

B) still images of IRI measurements in the absence and presence of 1.5 and 3 µM of mEos3.2-QAE .

C) quantitive IRI determination . Ice growth rates are determined as a function of concentration of QAE(T18N) concentration. The black line shows a sigmoidal nonlinear regression through the experimental values (black dots). $C_i$ represents the inflection point of the curve and indicates the activity of the AFP.

Scale bar: 10 µm

**Figure S4 corresponding to figure 2: representative sptPALM traces of ice-bound and coverslip immobilized mEos3.2-QAE shows highly comparable, immobile dynamics**

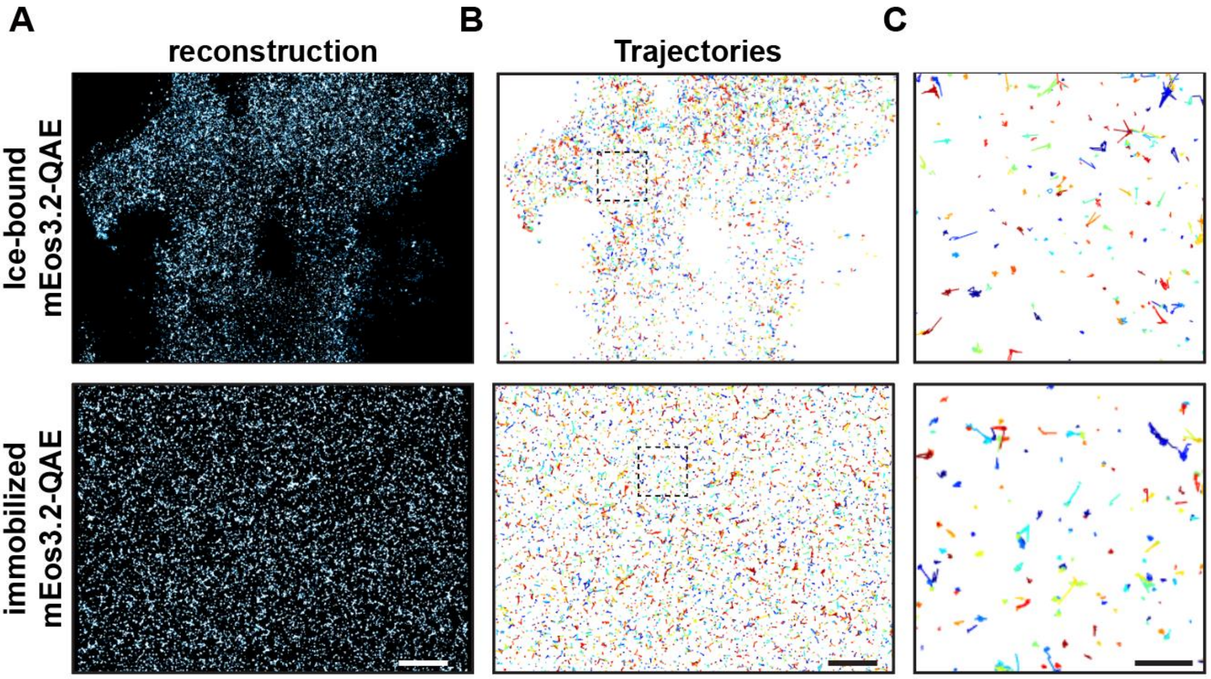


A) SMLM Reconstruction of ice-bound and coverslip immobilized mEos3.2-QAE at 3uM concentrations.
B) Tracks corresponding to acquisition in a of AFPs that were observed for a minimum of 5 frames. Individual tracks are randomly color coded.
C) Zoom of the tracks in the selected area in B
Scale bar: 5 µm (b), 1 µm (c)

**Figure S5 corresponding to figure 3: Activity of QAE(T18N) AFP upon fusion to fluorescent protein**

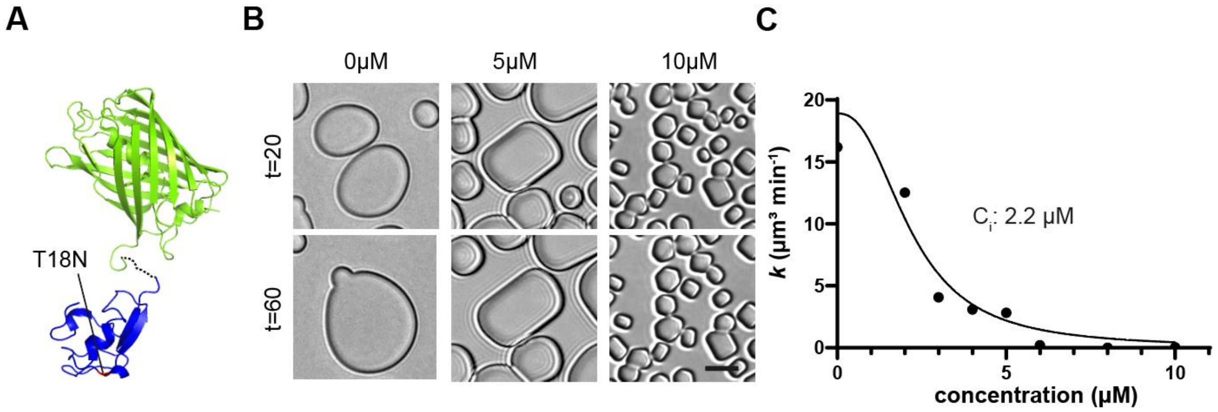


A) Structure of the mEos3.2-QAE(T18N) fusion. QAE AFP is depicted in blue, mEos in green and the linker in black dotted line. Threonine 18 is depicted in red.

B) still images of IRI measurements in the absence and presence of 5 and 10 μM of mEos3.2-QAE(T18N) .

C) Quantitive IRI determination . Ice growth rates are determined as a function of concentration of QAE(T18N) concentration. The black line shows a sigmoidal nonlinear regression through the experimental values (black dots). $C_i$ represents the inflection point of the curve and indicates the activity of the AFP.

Scale bar: 10 μm

### SUPPLEMENTAL TABLES

**Table S1: Protein sequences of the recombinant antifreeze proteins used in this study.**

| Construct | Amino Acid Sequence |
|---|---|
| mNeonGreen-**QAE**-6xHis<br><br>*Threonine 18 | MGVSKGEEDNMASLPATHELHIFGSINGVDFDMVGQGTGNPNDGYEELNLK<br>STKGDLQFSPWILVPHIGYGFHQYLPYPDGMSPFQAAMVDGSGYQVHRTMQ<br>FEDGASLTVNYRYTYEGSHIKGEAQVKGTGFPADGPVMTNSLTAADWCRS<br>KKTYPNDKTIISTFKWSYTTGNGKRYRSTARTTYTFAKPMAANYLKNQPMY<br>VFRKTELKHSKTELNFKEWQKAFTDVMGMDELYKEFGAGM**NQASVVANQ**<br>**LIPINTALTLVMMRSEVVTPVGIPAEDIPRLVSMQVNRAVPLGTTLMPD**<br>**MVKGYAA**LEHHHHHH* |
| mEos3.2-**QAE**-6xHis | MGSAIKPDMKIKLRMEGNVNGHHFVIDGDGTGKPFEGKQSMDLEVKEGGP<br>LPFAFDILTTAFHYGNRVFAKYPDNIQDYFKQSFPKGYSWERSLTFEDGGICN<br>ARNDITMEGDTFYNKVRFYGTNFPANGPVMQKKTLKWEPSTEKMYVRDGV<br>LTGDIEMALLLEGNAHYRCDFRTTYKAKEKGVKLPGAHFVDHCIEILSHDK<br>DYNKVKLYEHAVAHSGLPDNARREFGAGM**NQASVVANQLIPINTALTLV**<br>**MMRSEVVTPVGIPAEDIPRLVSMQVNRAVPLGTTLMPDMVKGYAA**LEH<br>HHHHH* |
| Halo-**QAE**-6xHis | MGAEIGTGFPFDPHYVEVLGERMHYVDVGPRDGTPVLFLHGNPTSSYVWRN<br>IIPHVAPTHRCIAPDLIGMGKSDKPDLGYFFDDHVRFMDAFIEALGLEEVVLV<br>IHDWGSALGFHWAKRNPERVKGIAFMEFIRPIPTWDEWPEFARETFQAFRTT<br>DVGRKLIIDQNVFIEGTLPMGVVRPLTEVEMDHYREPFLNPVDREPLWRFPN<br>ELPIAGEPANIVALVEEYMDWLHQSPVPKLLFWGTPGVLIPPAEAARLAKSL<br>PNCKAVDIGPGLNLLQEDNPDLIGSEIARWLSTLEISGEFGAGM**NQASVVAN**<br>**QLIPINTALTLVMMRSEVVTPVGIPAEDIPRLVSMQVNRAVPLGTTLMPD**<br>**MVKGYAA**LEHHHHHH* |
| Halo-**1wfa**-6xHis | MGAEIGTGFPFDPHYVEVLGERMHYVDVGPRDGTPVLFLHGNPTSSYVWRN<br>IIPHVAPTHRCIAPDLIGMGKSDKPDLGYFFDDHVRFMDAFIEALGLEEVVLV<br>IHDWGSALGFHWAKRNPERVKGIAFMEFIRPIPTWDEWPEFARETFQAFRTT<br>DVGRKLIIDQNVFIEGTLPMGVVRPLTEVEMDHYREPFLNPVDREPLWRFPN<br>ELPIAGEPANIVALVEEYMDWLHQSPVPKLLFWGTPGVLIPPAEAARLAKSL<br>PNCKAVDIGPGLNLLQEDNPDLIGSEIARWLSTLEISGEFGGGGS**DTASDAAA**<br>**AAALTAANAKAAAELTAANAAAAAAATARGAG**LEHHHHHH* |

**Table S2. Experimental conditions and imaging parameters for sptPALM acquisitions.**

| Figure | Ice orientation | protein | concentration | Exposure time (ms) |
|---|---|---|---|---|
| Fig 1B | Parallel | mEos3.2-QAE | 3.8 µM | 30 |
| Fig 2A-G | Parallel | mEos3.2-QAE | 1 µM | 15 |
| Fig 2H | Parallel | mEos3.2-QAE | 2.5 µM | 40 |
| Fig 2I,J | Parallel | mEos3.2-QAE | 1,3,10 µM | 40 |
| Fig 2L-O | perpendicular | mEos3.2-QAE | 2.5 µM | 40 |
| Fig 3E,F | perpendicular | mEos3.2-QAE; mEos3.2-QAE(T18N) | 10 µM | 40 |
| Fig 3H,I | perpendicular | Halo-QAE; Halo-QAE(T18N); Halo-wfAFP | 10,10,20 µM respectively | 20 |
| Fig S4 | Parallel | mEos3.2-QAE | 3µM | 40 |

### SUPPLEMENTAL MOVIES

**Movie S1:** tracking single QAE AFPs on polycrystalline ice by sptPALM. This movie corresponds to Figure 2A-G. sptPALM acquisition with 15ms interval of mEos3.2-QAE played. Raw acquisition (left) and corresponding tracks (right) are displayed. Total time 10.4 sec, scale bar 5 µm.

**Movie S2:** Different dynamics of Halo-tagged QAE, QAE(T18N) and wfAFP at the ice water interface. Movie corresponds to Figure 3H,I. paJF646 labelled AFPs were imaged on interfaces oriented perpendicular to the imaging plane. Total time 10 sec Scale bar 5 µm.

**Movie S3:** Zoomed in view of individual binding events at interfaces for Halo-tagged QAE, QAE(T18N) and wfAFP. Movie corresponds to Figure 3H,I. Raw acquisition with the superimposed interface (left) and corresponding tracks (right) are displayed. Total time 100 milliseconds, scale bar 2 µm.
